# ATP energy-independently modulates the folding equilibrium of ALS-causing C71G-hPFN1

**DOI:** 10.1101/2023.08.24.554696

**Authors:** Jian Kang, Liangzhong Lim, Jianxing Song

**Author notes:** J. K. and L.Z.L. contributed equally to this work. **Competing Interests:** No competing interests were disclosed.

## Abstract

C71G is the most aggregation-prone and toxic mutant of 140-residue human profin-1 (hPFN1) that causes familial ALS by gain of toxicity, but its underlying mechanisms still remain unknown. C71G-hPFN1 exists in an equilibrium between folded and unfolded states, whose dynamic/thermodynamic properties and modulation are not yet defined. Here, we utilized NMR to quantify the populations to be 55.2% and 44.8% respectively for folded and unfolded states exchanging at 11.7 Hz. Intriguingly, even the folded state of C71G-hPFN1 has increased ps-ns flexibility and reduced thermodynamic stability, thus rationalizing its high aggregation-proneness. Strikingly, C71G-hPFN1 provides a unique model to unambiguously visualize the effects of ATP and 11 related molecules on its folding equilibrium by NMR. Unexpectedly, ATP completely converted C71G-hPFN1 into the folded state at 1:2, which is physiologically relevant in most living cells. By contrast, TMAO, a well-known protein-folding inducer, showed no detectable conversion even at 1:2000. Surprisingly, the inducing capacity of ATP comes from its triphosphate group, but free triphosphate strongly triggered aggregation. The inducing capacity was determined to rank as: ATP = ATPP = PPP > ADP = AMP-PNP = AMP-PCP = PP, while AMP, Adenosine, P and NaCl showed no detectable capacity. Mechanistically, ATP and triphosphate appear to enhance the intrinsic folding capacity encoded by the sequence. Therefore, by joining Adenosine and triphosphate ATP appears to integrate three abilities: inducing folding, inhibiting aggregation and increasing stability. Our study provide a mechanism for the finding that some single-cell organisms still use polyphosphates as primordial chaperones. Moreover, ATP continue to play foundational roles in modern cells, shedding light on the longstanding enigma of the age-related onset of FALS, which coincides with the ageing-dependent reductions in ATP concentrations.

## Introduction

Many proteins need to fold from the unfolded state (U) into the folded state (F) for their functions (1-7). On the other hand, cell is extremely crowded in which protein concentrations can exceed 100 mg/ml, ∼3 mM of an ‘average’ protein (8). As the folded state of proteins is only marginally stable, some genetic mutations are sufficient to destabilize it, thus leading to their aggregation in cells, which is a pathological hallmark of aging and all neurodegenerative diseases (9-11). Modern cells handle protein folding and aggregation problems mainly with supramolecular machineries energetically driven by ATP (12), the universal energy currency which only requires micromolar concentrations for its previously-known functions (13). Mysteriously, however, ATP has very high concentrations in all living cells ranging from 2 to 12 mM dependent of cell types. For example, the vertebrate lens, a metabolically quiescent organ without mitochondrion, still maintains ATP concentrations of ∼7 mM (13-16).

Only recently, ATP has been decoded to energy-independently control protein hemostasis at concentrations >mM. In this context, ATP can behave as a biological hydrotrope to dissolve protein liquid-liquid phase separation (LLPS) and aggregates (15,16). We also found that ATP acts as a bivalent binder to induce and subsequently dissolve LLPS of intrinsically-disordered proteins (17-20) as well as to kinetically or/and thermodynamically inhibits amyloid fibrillation of well-folded domains (21,22). In particular, ATP appears to also act as a hydration mediator to antagonize the crowding-induced destabilization of the human eye-lens protein γS-crystallin without strong and specific binding (23). Nevertheless, so far no direct study has been reported regarding whether ATP can modulate protein folding equilibrium, the core event of protein hemostasis.

Amyotrophic lateral sclerosis (ALS) is the most common motor neuron disease which was first described in 1869 but its mechanism still remains a mystery (10). Most ALS cases are sporadic (90%) (SALS) whereas 10% are familial ALS (FALS). 140-residue human profilin-1(hPFN1) is a small protein that regulates actin polymerisation as well as other cellular functions. hPFN1 adopts a well-structured globular fold with a seven-stranded antiparallel β-sheet sandwiched by N- and C-terminal α-helices on one face of the sheet; and three small helical regions on the opposite face (24). Intriguingly, several mutations of hPFN1 have been identified to cause FALS but the exact mechanism remains unknown. Biophysically, these mutations have been shown to thermodynamically destabilize hPFN1 with the extent correlated to their tendency of aggregation *in vitro* and *in vivo* (25-27). Out of ALS-causing mutations on hPFN1, C71G with Cys71 mutated to Gly has been characterized to most severely destabilize the structure and consequently C71G-hPFN1 became highly prone to aggregation in buffers and consequently the attempts to determine its crystal structure has all failed (25-27). Most strikingly, C71G-hPFN1 has been demonstrated to be the most toxic and to cause ALS phenotypes in mice by gain of toxicity (28).

Previously, by an exhaustive screening, we have successfully identified the optimized buffer, thus allowing the high-resolution NMR characterization to unravel that C71G-hPFN1 co-exists between the unfolded and folded states characteristic of two sets of HSQC peaks (27). This observation suggests the existence of an energy barrier to separate the unfolded and folded states of C71G-hPFN1 (29,30). Remarkably, C71G-hPFN1 in folding equilibrium between two states may be established to be a unique model for directly visualizing the effect of small molecules on folding and aggregation of C71G-hPFN1 by NMR at the residue-resolution. On the other hand, so far, no measurement has been conducted to quantify the dynamic parameters of the conformational equilibrium such as the populations and exchange rate, as well as to define thermodynamic and dynamic stability of C71G-hPFN1. In particular, one question of both fundamental and therapeutic interest is whether ATP universally existing in all living cells with concentrations >mM can modulate this folding equilibrium.

In the present study, to shed light on the mystery of ALS caused by C71G-hPFN1, we first aimed to map out NMR dynamics of the conformational equilibrium on both ps-ns and μs-ms time scales, as well as thermodynamic stability of C71G-hPFN1. Subsequently, we assessed whether ATP is capable of modulating the folding equilibrium by directly visualizing the effects of ATP and 11 related molecules on the two states of C71G-hPFN1, which include ADP, AMP, Adenosine, ATPP, AMP-PNP, AMP-PCP, sodium triphosphate, sodium pyrophosphate and sodium phosphate, as well as sodium chloride and trimethylamine N-oxide (TMAO) (Fig. 1). The results not only led to the definition of the populations of the folded and unfolded states of C71G-hPFN1 as well as their exchange rate, but also unveiled that the ps-ns backbone flexibility of the folded state has largely increased as compared to those of WT-hPFN1, generally consistent with its considerably reduced thermodynamic stability. These results together provide a biophysical mechanism for the high aggregation-proneness of C71G-hPFN1. Most strikingly, the study has decoded for the first time that without any strong and specific binding, ATP could completely convert the unfolded state into the folded state even at a ratio of 1:2 (C71G:ATP). By contrast, TMAO, currently the best-known molecule with the general capacity in inducing protein folding, failed to show any detectable conversion even at a ratio up to 1:2000 at which TMAO had extensive interactions with C71G-hPFN1 as well as induced its precipitation. Most unexpectedly, the inducing capacity of ATP has been identified to come from its triphosphate group. However, although the isolated triphosphate (PPP) was able to induce the folding as effectively as ATP, it showed strong ability to trigger protein aggregation. On the other hand, ADP needed much higher ratio (1:8) than ATP to achieve the complete conversion while AMP no longer has the detectable capacity even at 1:400. Similarly, the isolated pyrophosphate has ADP-like inducing capacity but again has strong ability to induce aggregation while phosphate only induced aggregation. Results together imply that only by joining Adenosine with triphosphate but not tetraphosphate or methylenediphosphonate or imidodiphosphate, nature marvelously engineers ATP which owns the integrated three abilities: to effectively induce protein folding, to inhibit aggregation and to enhance thermodynamic stability. Our results also provide a possible mechanism for the previous finding that in some single-cell organisms, inorganic polyphosphates can function as a primordial chaperone (31,32). The capacity of ATP in inducing protein folding at physiologically-relevant concentrations implies that ATP may play a previously-unknown role in preventing the aggregation of disease-causing mutants, thus explaining that the individual carrying the ALS-causing mutants has ALS onset only after a certain age likely when ATP concentration reduced to a certain level.

**Fig. 1.**
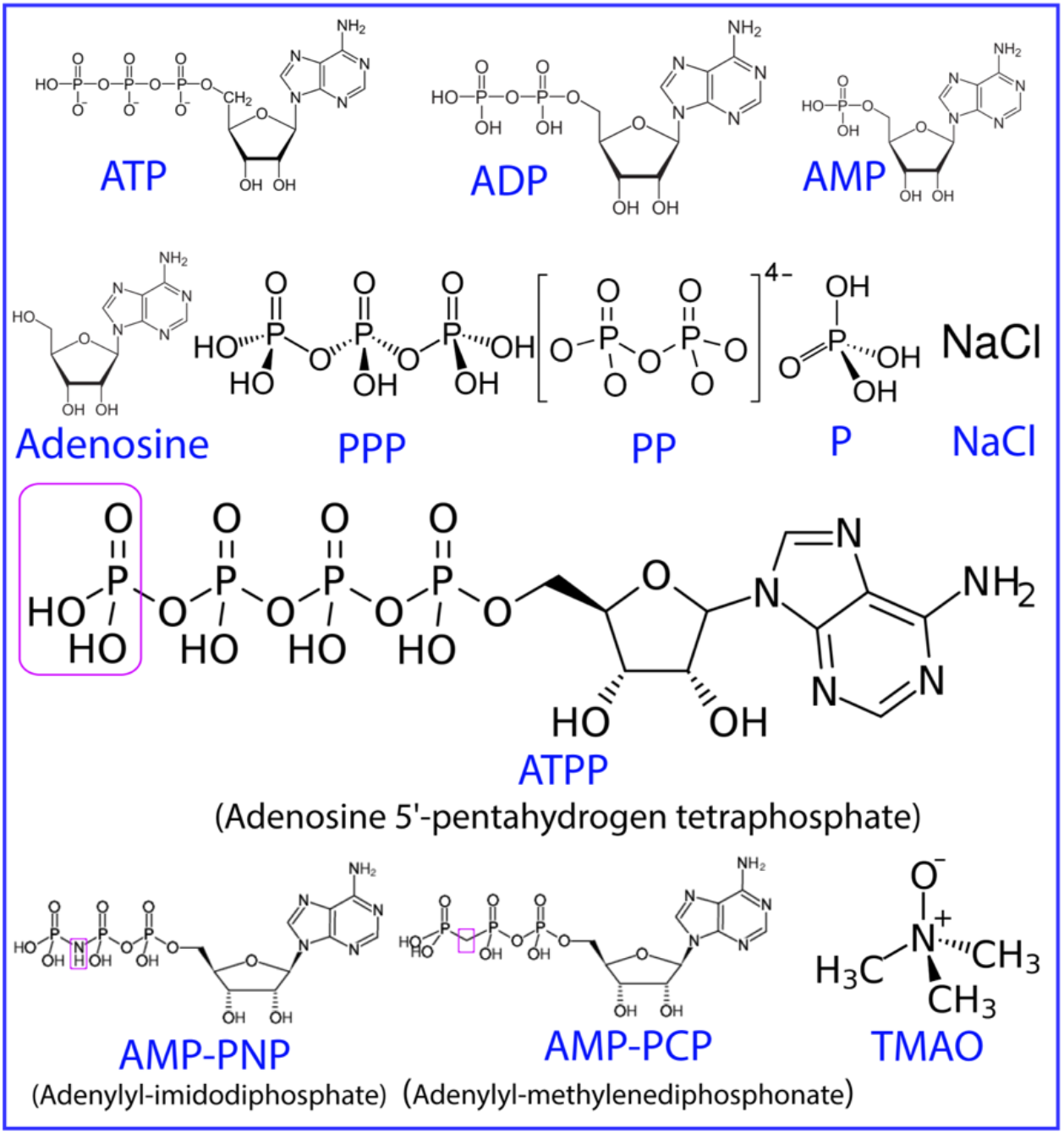
ATP and 11 related compounds utilized in the present study.

## Results

### NMR quantification of the conformational equilibrium of C71G-hPFN1

Previously, by exhaustively screening buffer conditions including pH, types and concentrations of salts, we have successfully identified the optimized buffer: 1 mM phosphate buffer at pH 6.0, in which the aggregation could be minimized and C71G-hPFN1 could be characterized by high-resolution NMR (27). In the present study, the low buffer salt concentration at 1 mM is also required to maximize the manifestation of the effects of ATP and related molecules.

As previously established, C71G-hPFN1 has a co-existence of the folded and unfolded states which manifest two sets of HSQC peaks: one from the folded state and the other from the unfolded state (27). To allow high-resolution NMR visualization here, by collecting and analyzing a series of triple-resonance 3D NMR spectra collected on ^15^N-/^13^C-labeled WT-hPFN1 and C71G-hPFN1 samples, we achieved sequential assignments of both WT-hPFN1 and C71G-hPFN1 except for some missing or overlapping peaks. As indicated by Fig. S1A, WT-hPFN1 and the folded state of C71G-hPFN1 have highly similar (ΔCα-ΔCβ) values, indicating that they have very similar secondary structures. By contrast, the absolute values of (ΔCα-ΔCβ) of the unfolded state are much smaller than those of its folded state (Fig. S1B), clearly suggesting that the unfolded state is highly disordered without any stable secondary structures (33).

In order to allow the comparison of ps-ns backbone dynamics between WT-hPFN1 and the folded state of C71G-hPFN1, here we also collected ^15^N backbone relaxation data T1, T2 and hNOE for both WT-hPFN1 and C71G-hPFN1 in the same buffer (Fig. S2). As shown in Fig. S2, most residues of WT-hPFN1 have very large hNOE, with the average of 0.82, typical of a well-folded protein (34-37). Subsequently we performed the “model-free” analysis, which generates squared generalized order parameters, S^2^, reflecting the conformational rigidity on ps-ns time scale. S^2^ values range from 0 for high internal motion to 1 for completely restricted motion in a molecular reference frame (34-37). As shown in Fig 2, the majority of the WT-hPFN1 residues has S^2^ > 0.76 (with the average value of 0.89), suggesting that WT-hPFN1 has very high conformational rigidity.

**Fig. 2.**
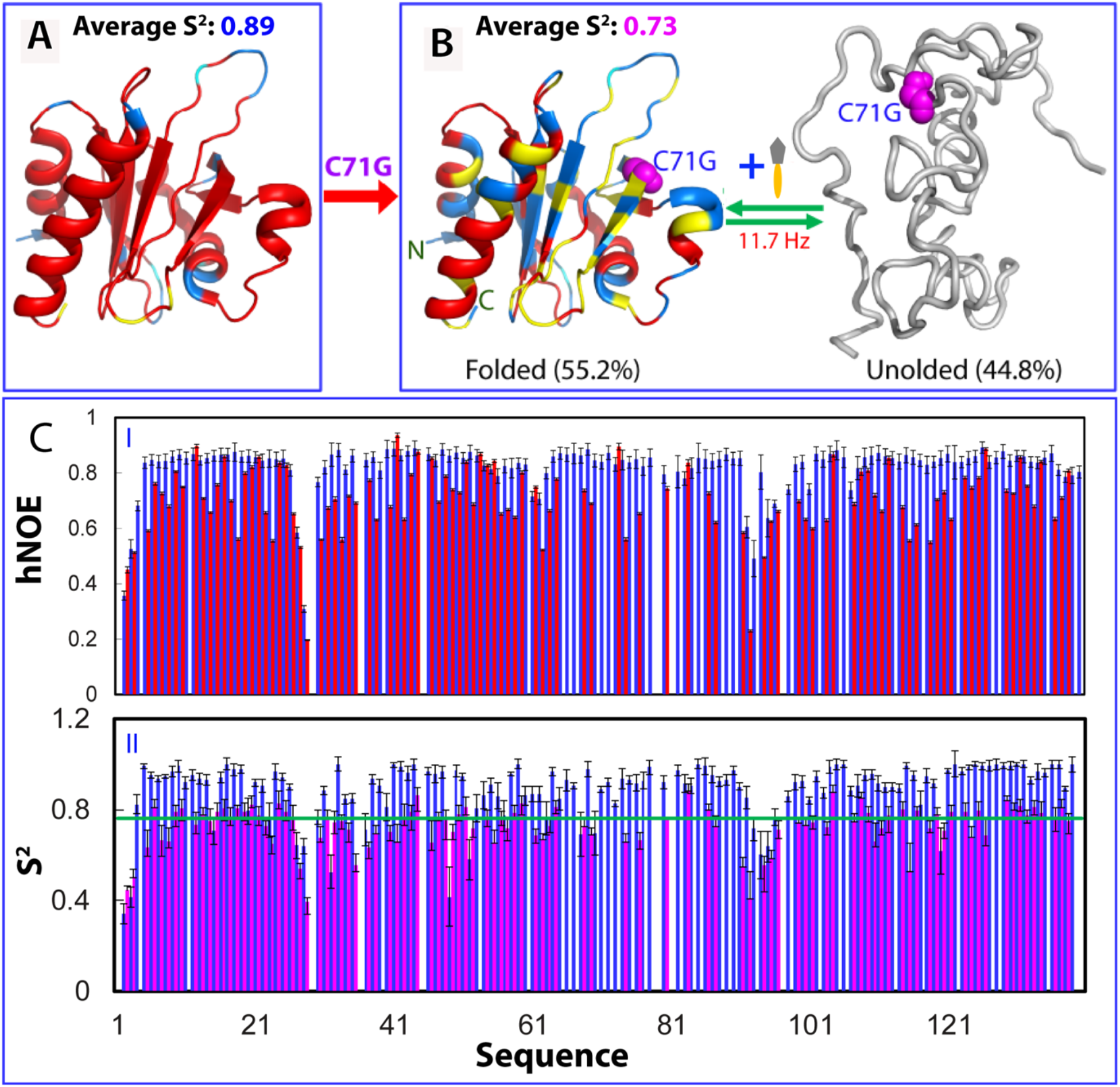
NMR quantification of ps-ns and μs-ms dynamics of the folding equilibrium. (A) The structure of WT-hPFN1 (PDB ID of 2PAV) with S^2^ values of WT-hPFN1 (the average value of 0.89) mapped on. (B) A diagram to show the co-existence of the folded (the average S^2^ value of 0.73) and unfolded states of C71G-hPFN1. The populations of the folded and unfolded states have been calculated to be respectively 55.2% and 44.8% which undergo a conformational exchange at 11.7 Hz. (C) {^1^H}-^15^N steady-state NOE intensities (I) and generalized squared order parameter (S^2^) (II) of WT-hPFN1 (blue) and the folded state of C71G-hPFN1 (red). The green line has a S^2^ value of 0.76 (average – STD of WT-hPFN1). S^2^ values are also mapped onto the folded state of C71G-hPFN1 in (B). Cyan is used for indicating Pro residues, and the yellow for residues with missing or overlapping HSQC peaks. Red is for residues with S^2^ values > 0.76 and blue for residues with S^2^ values < 0.76.

As the folded and unfolded states have very different relaxation behaviors, it is impossible to derive their populations from the intensity of HSQC peaks. Therefore, to quantify the populations and exchange parameters for the folding equilibrium of C71G-hPFN1, we have collected and then analyzed HSQC-NOESY spectrum of C71G-hPFN1. Strikingly, the cross peaks resulting from the exchange of the folded and unfolded states have been observed (Fig. S3). This allowed the successful calculation of the populations and exchange rate for the unfolded and folded states of C71G-hPFN1 by using the well-established NMR methods (29,30). As presented in Table S1, C71G-hPFN1 has the populations of 55.2% and 44.8% respectively for the folded and unfolded states with an exchange rate of ∼11.7 Hz (∼85.5 milli-second) (Fig. 1B).

We also analyed the relaxation data for C71G-hPFN1 (Fig. S2). As shown in I of Fig. 1C, C71G-hPFN1 have the average hNOE value of 0.70 with most residues having hNOE values smaller than those of WT-hPFN1, indicating that the folded state of C71G-hPFN1 has backbone dynamics higher than those of WT-hPFN1 on ps-ns time scale (34-37). To facilitate the comparison, we also performed the “model-free” analysis. As shown in II of Fig 1C, many residues of the folded state of C71G-hPFN1 have S^2^ < 0.76 (Fig. 1A) (with the average value of 0.73), revealing that even the folded state of C71G-hPFN1 becomes more flexible than WT-hPFN1 on ps-ns time scale.

Furthermore, the overall rotational correlation times (τc) were calculated to be 7.5 ns and 7.8 ns respectively for WT-hPFN1 and C71G-hPFN1. The results imply that C71G-hPFN1 becomes slightly less compact. To conform this, here by use of pulsed field gradient NMR self-diffusion measurements (38), we also measured their translational diffusion coefficients to be 1.12 ± 0.03 × 10^−10^ m^2^/s for WT-hPFN1, and 1.03 ± 0.02 × 10^−10^ m^2^/s for C71G-hPFN1, which independently indicate that the folded state of C71G-hPFN1 indeed becomes slightly less compact.

### ATP induces folding of the unfolded state of C71G-hPFN1

Here, we first assessed whether ATP has any specific binding pocket on WT-hPFN1 by titrating ATP into the WT-PFN1 sample as monitored by NMR. As shown in I of Fig. S4, even with the molar ratio up to 1:400 (hPFN1:ATP), ATP induced no large shift of HSQC peaks of WT-PFN1 except for those of His120 and Gly121. As His120 and Gly121 are located on an exposed loop (II of Fig. S4), it is most likely that the shift of their HSQC peaks is resulting from non-specific electrostatic effect from the highly negatively-charged ATP. The result suggests that unlike the folded nucleic-acid-binding domains (17) such as TDP-43 NTD (30) and FUS RRM (21) with specific ATP-binding pockets constituted by many residues which were all significantly perturbed by ATP titrations, WT-hPFN1 without any known activity in binding nucleic acid also has no specific binding pocket for ATP.

Strikingly, when ATP was added into the C71G-hPFN1 sample even only at a molar ratio of 1:0.5 (C71G:ATP), the intensity of HSQC peaks of the unfolded state became reduced while those of the folded state slightly increased (Fig. S5 and S6). The 1D peak intensity of the methyl group from the unfolded state also became reduced while those of the folded state slightly increased (III of Fig. S5). When ATP was added to 1:1, the HSQC peak intensity of the unfolded state became further reduced while those of the folded state increased (Fig. 3 and S5). At 1:2, the 1D peak intensity of the methyl group of the folded state further increased (Fig. 3B) while HSQC peaks of the unfolded state became completely disappeared (Fig. 3C). Further addition of ATP to 1:20 (1 mM) only induced further shifts of some HSQC peaks of the unfolded state (IV of Fig S4). These results indicate that even at 1:2, ATP is capable of completely shifting the conformational equilibrium to the folded state. Interestingly, no considerable shift was observed for HSQC peaks of the folded state of C71G-hPFN1 except for those of His120 and Gly121 (Fig. 3C). This clearly indicates that like WT-hPFN1 (Fig. S4), C71G-hPFN1 also has no specific binding pocket for ATP. We also increased the C71G-hPFN1 concentration to 100 μM, and the ATP concentration required to completely convert the unfolded state also doubled (200 μM). This result suggests that the conversion by ATP is dependent of the molar ratio between C71G-hPFN1 and ATP.

**Fig. 3.**
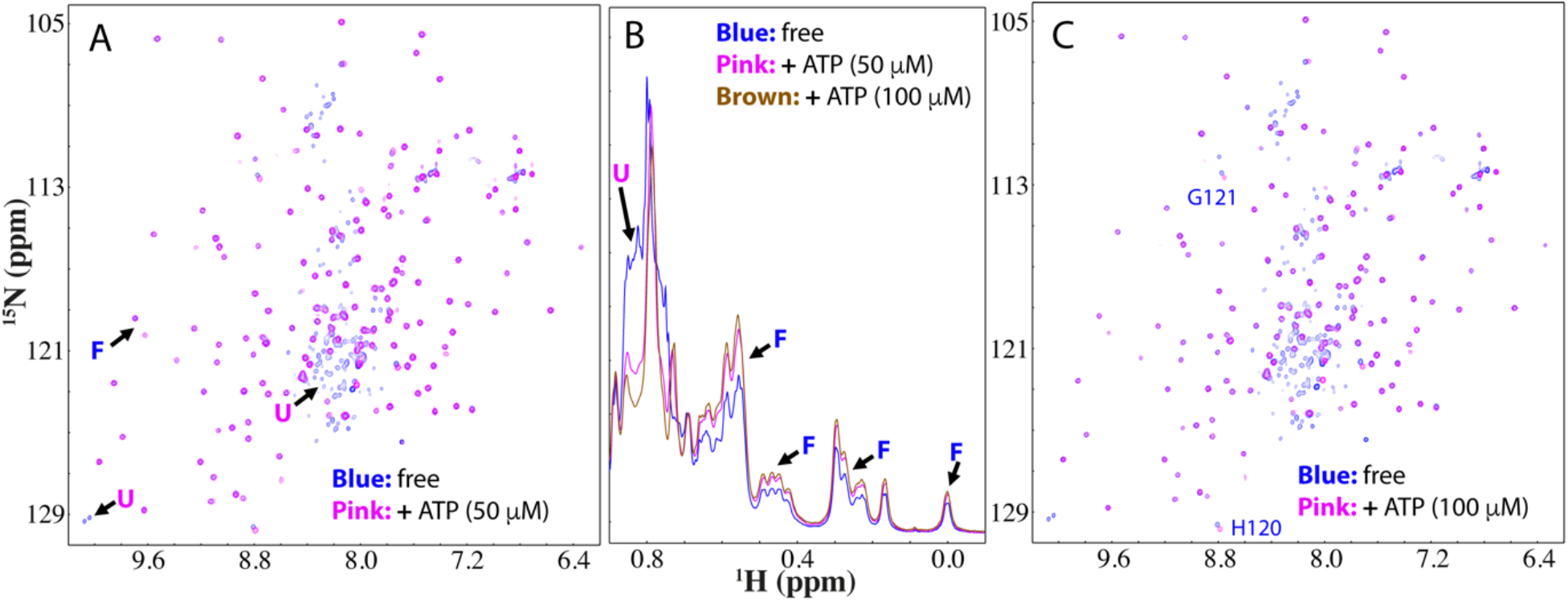
ATP completely converted the unfolded state into the folded state. **(A)**Superimposition of HSQC spectra of ^15^N-labeled C71G-hPFN1 at a concentration of 50 μM in the absence (blue) or in the presence of ATP (pink) at a molar ratio of 1:1. Some characteristic NMR signals of the folded (F) and unfolded (U) states were indicated by arrows(B)Up-field 1D NMR spectra of C71G-hPFN1 in the presence of ATP at different ratios. (C) Superimposition of HSQC spectra of C71G-hPFN1 in the absence (blue) and in the presence of ATP (pink) at a molar ratio of 1:2. Two residues (His120 and Gly121) of the folded state with shifted HSQC peaks are labelled.

### ADP, AMP and Adenosine have differential inducing capacity

To determine the group and mechanism of ATP molecule responsible for this inducing capacity, we systematically titrated C71G-hPFN1 with a list of 11 related molecules (Fig. 1). For ADP, only at 1:8 (C71G:ADP), HSQC peaks of the unfolded state became completely converted into the folded state. Strikingly, HSQC peaks with ADP at 1:8 are highly superimposable to those with ATP at 1:2 (Fig. S7). This indicates that ADP still has the capacity in inducing the folding of the unfolded state but the capacity is much weaker than that of ATP.

We also titrated with AMP (Fig. S8), but even with the concentrations up to 20 mM (1:400), AMP was still unable to convert the unfolded state into the folded state. We further titrated with Adenosine and even with the highest concentrations of 5 mM due to its low solubility, no significant change was detected (Fig. S9). The results strongly imply that the capacity of ATP in inducing the folding might come from its triphosphate group.

### The inducing capacity of ATP comes from its triphosphate group

To define the roles of their phosphate groups of ATP, ADP and AMP in inducing the folding, we then systematically titrated C71G-hPFN1 with sodium salts of triphosphate (PPP), pyrophosphate (PP) and phosphate (P). For PPP, at 1:1 (C71G:PPP), the HSQC peak intensity of the unfolded state also became significantly reduced (Fig. 4 and S10). Furthermore, the 1D peak intensity of the methyl group of the unfolded state also became reduced while those of the folded state slightly increased (Fig 4B). At 1:2, HSQC peaks of the unfolded state became completely disappeared (Fig. 4C), and the intensity of the methyl group of the folded state further increased (Fig. 4B). Very strikingly, C71G-hPFN1 with PPP at 1:2 has both HSQC (Fig. 4D) and 1D (Fig. 4B) spectra very similar to those with ATP at 1:2. The result suggests that ATP and PPP induce the folding of the unfolded state of C71G-hPFN1 with the highly-similar effectiveness and mechanism.

**Fig. 4.**
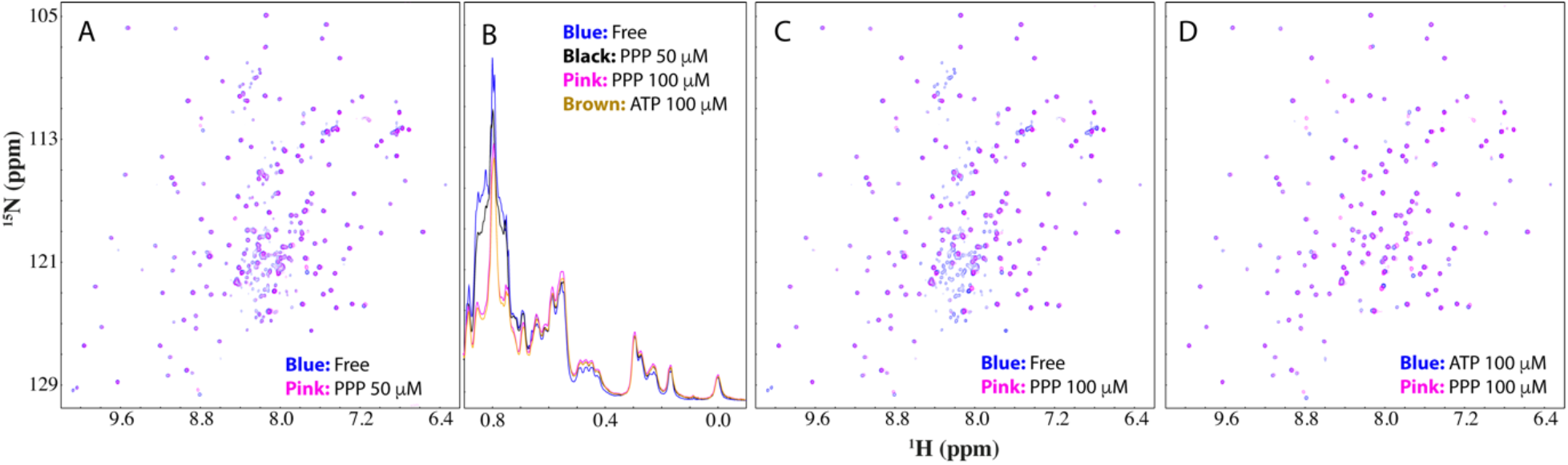
Triphosphate (PPP) converted the unfolded state into the folded state. Superimposition of HSQC spectra of ^15^N-labeled C71G-hPFN1 at a concentration of 50 μM in the absence (blue) and in the presence of PPP (pink) at a molar ratio of 1:1 (A), and 1:2 (C).(B) Up-field 1D NMR spectra of C71G-hPFN1 in the presence of PPP at different ratios. (D) Superimposition of HSQC spectra of ^15^N-labeled C71G-hPFN1 in the presence of ATP at 1:2 (blue) or PPP (pink) at 1:2.

On the other hand, addition of PPP at 1:4 resulted in the broadening of NMR signals and after one hour the sample started to form white precipitates and NMR signals become too weak to be detectable. The results revealed that although the capacity of ATP in inducing the folding comes from its triphosphate group, the free triphosphate also owns a very strong ability to induce severe aggregation of C71G-hPFN1. Mechanistically, as we previously showed (11), highly disordered and partially-folded proteins have many exposed hydrophobic patches Consequently, they will be driven to aggregate by hydrophobic interactions upon contacting charged molecules which can impose electrostatic screening effects (11). Here as revealed by the above NMR studies on both ps-ns and μs-ms dynamics, not only the unfolded state of C71G-hPFN1 is highly disordered, its folded state also has an increased ns-ps dynamics. As such, even though the unfolded state of C71G-hPFN1 was completely converted into the folded state by PPP, the folded state of C71G-hPFN1 still has mutation-causing packing defects and dynamically exposed hydrophobic patches, which will be unavoidably triggered to aggregate in the presence of the highly negatively charged PPP. By contrast, the aromatic base ring of ATP might be dynamically clustered onto the exposed hydrophobic patches of C71G-hPFN1 to shield the screening effects imposed by triphosphate group to some extent, thus attenuating the aggregation induced by PPP.

Indeed, we further titrated with pyrophosphate (PP). As shown in Fig. S11, at 1:6 (C71G:PP), the HSQC peaks of the unfolded state still retained despite the reduction of their intensity. At 1:8, although the HSQC peaks of the unfolded state disappeared, the intensity of 1D and HSQC peaks of the folded state also reduced. In particular, after about one hour, the visible precipitates could be observed and NMR signals become too weak to be detectable.

Subsequently, we carried out titrations with sodium phosphate (Fig. S12) and chloride (Fig. S13), and both failed to convert the unfolded state into the folded state with the concentration up to 5 mM (1:100) for sodium phosphate (Fig. S12), and 10 mM (1:200) for NaCl (Fig. S13), where NMR samples started to show visible precipitation after one hour.

These results with PPP, PP, P and NaCl strongly imply that the inducing capacity of PPP and PP is not mainly due to their high ionic strength because if assume that all are sodium salts, the ionic strength of sodium triphosphate (PPP) and sodium pyrophosphate (PP) are maximally 15-time and 10-time stronger than sodium chloride while only 2.5-time and ∼1.7-time stronger than sodium phosphate (39). Nevertheless, sodium chloride at 10 mM with an ionic strength much larger than those of PPP at 0.1 mM and PP at 0.4 mM showed no capacity at all in inducing the folding of the unfolded state but only triggered the aggregation of C71G-hPFN1.

### The inducing capacity depends on the atoms linking the beta and gamma phosphates

We further assessed the effects on the folding equilibrium of three nonnatural ATP analogues, namely Adenosine 5’-(pentahydrogen tetraphosphate) (ATPP), Adenylyl-imidodiphosphate (AMP-PNP) and Methyleneadenosine 5-triphosphate (AMP-PCP) (Fig. 1).Very unexpectedly, ATPP induced the complete conversion of the unfolded state into the folded state at 1:2 (C71G:ATPP) which is very similar to ATP (Fig. S14A). This result indicates that the inclusion of an extra phosphate failed to increase the capacity of ATP to induce folding. By contrast, ATPP suddenly showed much higher ability than ATP to trigger the aggregation of the C71G-hPFN1 protein as evidenced by the result at 1:4 (C71G:ATPP), the C71G protein started to aggregate and many HSQC peaks became too broad to be detected. Further addition of ATPP immediately triggered visible precipitation and no NMR signal could be detected.

Furthermore, upon replacing the oxygen atom linking the beta and gamma phosphates with carbon atom, the capacity of AMP-PCP (Fig. 1) to induce folding reduced to the level very similar to that of ADP: only at a ratio of 1:8 (C71G:AMP-PCP) the unfolded state was completely converted into the folded state (Fig. S14B). On the other hand, unlike ADP which triggered no precipitation even at 20 mM, upon adding AMP-PCP with concentrations at 1 mM, the C71G-hPFN1 protein also started to aggregate and precipitate after one hour.

Intriguingly, AMP-PNP with the oxygen atom replaced by nitrogen atom also has the capacity in inducing folding very similar to that of ADP and AMP-PNP (Fig. S14C). However, its ability to trigger aggregation is even higher than that of AMP-PCP: even at 1:10 (C71G:AMP-PNP) with the AMP-PNP concentration of 0.5 mM, the C71G-hPFN1 protein started to precipitate after one hour.

### TMAO failed to shift the equilibrium but only triggered aggregation

Currently, the best-known molecule with the general capacity in inducing protein folding is the natural osmolyte, trimethylamine N-oxide (TMAO) (3,40-42), which is generated from choline, betaine, and carnitine by gut microbial metabolism (42). Here, we titrated TMAO into the C71G-hPFN1 sample (Fig. 5). At 1:2 to 1:20 (C61G:TMAO), except for the shift of several peaks, no large change were observed for HSQC peaks of both folded and unfolded states. At 1:200, extensive shifts were observed for HSQC peaks of both folded and unfolded states. However, even at 1:2000, HSQC peaks of both folded and unfolded states still co-existed but the intensity of 1D peaks for both states reduced considerably. At 1:2000, the C71G-hPFN1 sample become completely precipitated after one hour and NMR signals became disappeared. The results clearly revealed that TMAO was unable to induce the folding of the unfolded state of C71G-hPFN1 even with the ratio up to 1:2000 at which C71G-hPFN1 started to precipitate. This is ib general consistent with previous reports that TMAO concentrations needed to reach ∼M to induce the folding of proteins (3,40,41).

**Fig. 5.**
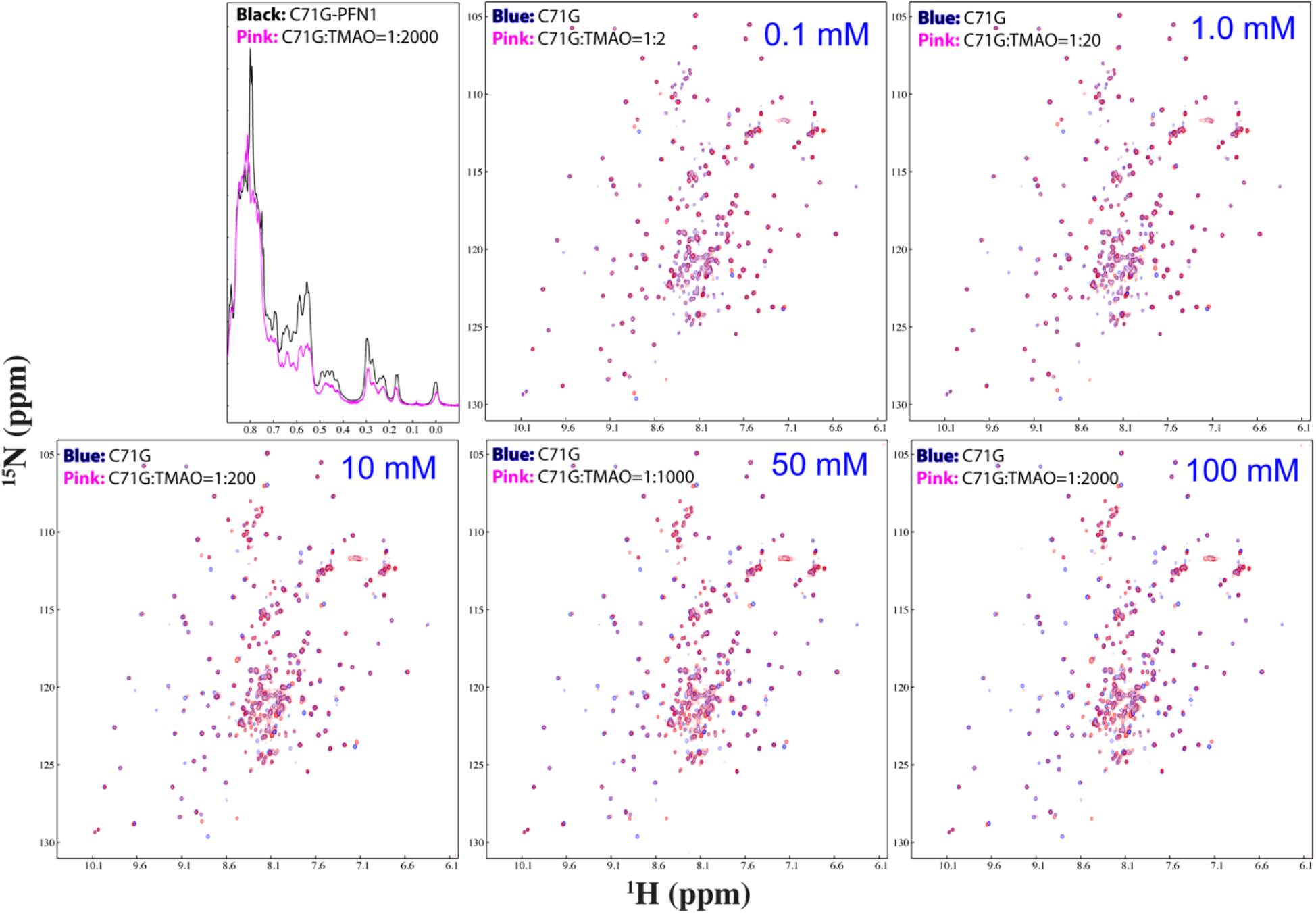
TMAO showed no capacity in converting the unfolded state into the folded state. Up-field 1D and HSQC spectra of C71G-hPFN1 at a concentration of 50 μM in the presence of TMAO at 1:2, 1:20, 1:200, 1:1000 and 1:2000 (C71G:TMAO) above which TMAO triggered a complete precipitation. For clarity, only 1D spectra of C71G-hPFN1 in the free state (black) and in the presence of TMAO at 1:2000 were presented.

### ATP enhances thermodynamic stability of the folded state of C71G-but not WT-hPFN1

We also measured the thermodynamic stability by differential scanning fluorimetric (DSF) method (21-23,43). Interestingly, WT-hPFN1 has a melting temperature (Tm) of 56 degree while the addition of ATP even up to 20 mM triggered no significant change (Fig. 5A). By contrast, C71G-hPFN1 at 10 μM without ATP has no cooperative unfolding signal (Fig. 5), likely due to the absence of tight tertiary packing or/and co-existence of two states (30,44,45). Interestingly in the presence of ATP of 20 μM (a molar ratio of 1:2), a cooperative unfolding signal was observed with Tm of 32 degree. Addition of ATP to 1 mM (1:100) led to the increase of Tm to 38 degree, and further increase to 40 degree at 20 mM (Fig. 5A).

By contrast, we also performed many DSF measurements of C71G-hPFN1 with addition of PPP at various ratios and only obtained unfolding curves with no cooperative unfolding signals but high noises, implying that although ATP and PPP both have similar capacity in inducing folding, PPP failed to enhance tight tertiary packing of the folded state or/and triggered dynamic aggregation even before the visible precipitation. Indeed, previously we found that a small protein could still have native-like NMR spectra although its tight packing was disrupted to different degrees (44,45).

## Discussion

Out of ALS-causing hPFN1 mutants identified so far, C71G-hPFN1 is the most toxic but the attempts to determine its crystal structure has all failed. Our previous NMR studies revealed that C71G-hPFN1 co-exists between the folded and unfolded states (27), but their conformations, dynamics, stability and equilibrium parameters remain to be defined. In the present study, we have successfully quantify the populations of the folded and unfolded states to be respectively 55.2% and 44.8%, which are exchanging at 11.7 Hz (∼85.5 milli-second). Moreover, although the folded state of C71G-hPFN1 has a similar conformation as WT-hPFN1, it has an increased backbone flexibility on ps-ns time scale, while its thermodynamic stability with Tm of 32 degree is also much lower than that of WT-hPFN1 (Tm of 56 degree). These results provide a biophysical mechanism for the observation that C71G-hPFN1 is highly aggregation-prone both *in vitr*o and *in vivo*. Briefly, due to the relatively small energy barrier separating the folded and unfolded states as reflected by the exchange rate of 11.7 Hz, the aggregation of the unfolded state will lead to the rapid conversion of the folded state into the unfolded state and consequently all C71G-hPFN1 protein finally become aggregated. Furthermore, even the folded state of C71G-hPFN1 may directly get aggregated due to its low dynamic and thermodynamic stability.

The co-existence of the folded and unfolded states with well-defined equilibrium parameters established C71G-hPFN1 composed of both α-helices and β-sheets without any disulfide bridge to be a unique model to unambiguously determine the effects of small molecules on folding by NMR visualizing the equilibrium shift of two C71G-hPFN1 states. Currently although there is a list of well-established model proteins with >100 residues for folding studies, to the best our knowledge, there is no protein with the co-existence in ab equilibrium between the folded and unfolded states which could be characterized by high-resolution NMR. Furthermore, for the helical proteins such as Myoglobin, they are very stable and thus it is challenging to identify the point mutation to destabilize them into the co-existence of the two states. On the other hand, multiple mutations will fundamentally alter the sequence-structure relationship and thus the multi-mutated sequences may become unfoldable. For other proteins rich in β-sheets such as Ribonucleases, alpha-Lactalbumin and Lysozyme, they contain >1 disulfide bridges which radically complicate the sequence-structure relationship. For example, even the partial removal of the disulfide bridges of the alpha-lactalbumin led to being trapped in molten globule whose NMR signals are too broad to be detected (46). Indeed, in the accompanying manuscript, we further studied 153-residue human superoxide dismutase 1 (hSOD1) with the Greek-key β-barrel fold stabilized by only 1 disulfide bridge (47). Its nascent form without the disulfide bridge as well as cofactors zinc and copper ions is completely unfolded, but could, however, be induced to reach an equilibrium between the folded and unfolded states specifically by zinc ion at 1:20 as well as generally by ATP at 1:8. Nevertheless, without the formation of the disulfide bridge, zinc ion or/and ATP could not completely convert the unfolded state into the folded state, which are separated by the energy barrier larger than that for the two states of C71G-hPFN1. This suggests that disulfide bridge is a key sequence factor which specifies the multi-step folding of hSOD1 (47). Most importantly, C71G-hPFN1 is the most toxic ALS-causing mutant and in this context the identification of any small molecules capable of inducing its folding holds fundamental and therapeutic implications for our better understanding and treatment of ALS pathogenesis.

Indeed, for the first time in the present study, we discovered that ATP has an energy-dependent capacity in completely converting the unfolded state into the folded state of C71G-hPFN1 even at 1:2, a very low ratio which can be satisfied in most, if not all living cells. By contrast, even with the ratio reaching 1:2000 at which extensive binding was observed and C71G-hPFN1 already started to precipitate, TMAO failed to show a detectable conversion. Very unexpectedly, by a systematic titration of 10 related molecules, the inducing capacity of ATP has been decoded to come from triphosphate which was previously proposed to act as a key intermediate for various prebiotic chemical reactions to generate building units for constructing primitive cells, thus allowing the Origin of Life (48,49).

The inducing capacity of these molecules was further determined here to rank as: ATP = ATPP = PPP > ADP = AMP-PNP = AMP-PCP = PP, while AMP, Adenosine, P and NaCl showed no detectable capacity. Intriguingly, ATPP, PPP, AMP-PNP, AMP-PCP, PP, P and NaCl all showed strong electrostatic screening effects to trigger the aggregation of C71G-hPFN1. The results indicate that the inducing capacity depends on the number of phosphate groups linked to Adenosine but triphosphate group in ATP has the maximal capacity as ATPP has no increased capacity. On the other hand, the atoms linking phosphoric atoms are also important as AMP-PNP and AMP-PCP with linking oxygen atom replaced respectively by nitrogen and carbon atoms all led to the reduced capacity as well as increased screening effect. Therefore, Adenosine in ATP and ADP appears to be able to shield the screening effects of PPP and PP groups. The results together imply that the linkage of Adenosine and triphosphate represents the best combination not only to maximize the capacity in inducing folding, but also to minimize the ability of triphosphate to trigger aggregation.

So, what could be the mechanism for ATP and triphosphate to induce the complete conversion of the unfolded state into the folded state of C71G-hPFN1 at 1:2? At such a low ratio, ATP has no strong and specific interaction with the folded state of C71G-hPFN1 and also no volume excluding effect is anticipated to operate. After exhaustively searching for the literatures on the mechanisms of protein folding, we believe that the most relevant is that ATP or triphosphate act to induce folding by interacting with protein hydration. Previously, it was proposed that the hydrogen bonding with water molecules or/and solvation, particularly for the protein backbone atoms play a key role in protein folding (2-4). The unfolded state (U) will be favored if the backbone atoms are highly hydrogen-bonded with water molecules, while the folded state (F) will be favored if the backbone atoms are involved in forming intramolecular hydrogen-bonds (3).

Here as illustrated by Fig. 6B, we propose here that ATP or triphosphate induces the folding by effectively interacting with water molecules hydrogen-bonded with the backbone atoms of the unfolded state and thus attracting water molecules out from hydrogen-bonding or/and hydration shell with its backbone atoms. Consequently, the backbone atoms of the unfolded state will be shifted to forming the intramolecular hydrogen bonds to favoring folding. Furthermore, most likely by using the aromatic ring of adenosine to be dynamically clustered over and thus protecting the hydrophobic patches from being exposed to bulk solvent, ATP further acquired a novel ability to enhance the thermodynamic stability of the folded state of C71G-hPFN1 with the defects in tertiary packing. Indeed, without ATP, C71G-hPFN1 had no cooperative thermal unfolding. In the presence of ATP at 20 μM (1:2), C71G-hPFN1 showed cooperative thermal unfolding with Tm of 32 degree, which increased to 40 degree in the presence of ATP at 20 mM (1:2000). By contrast, WT-hPFN1 without the defects in tertiary packing showed no change of Tm even with ATP at 20 mM (1:2000).

**Fig. 6.**
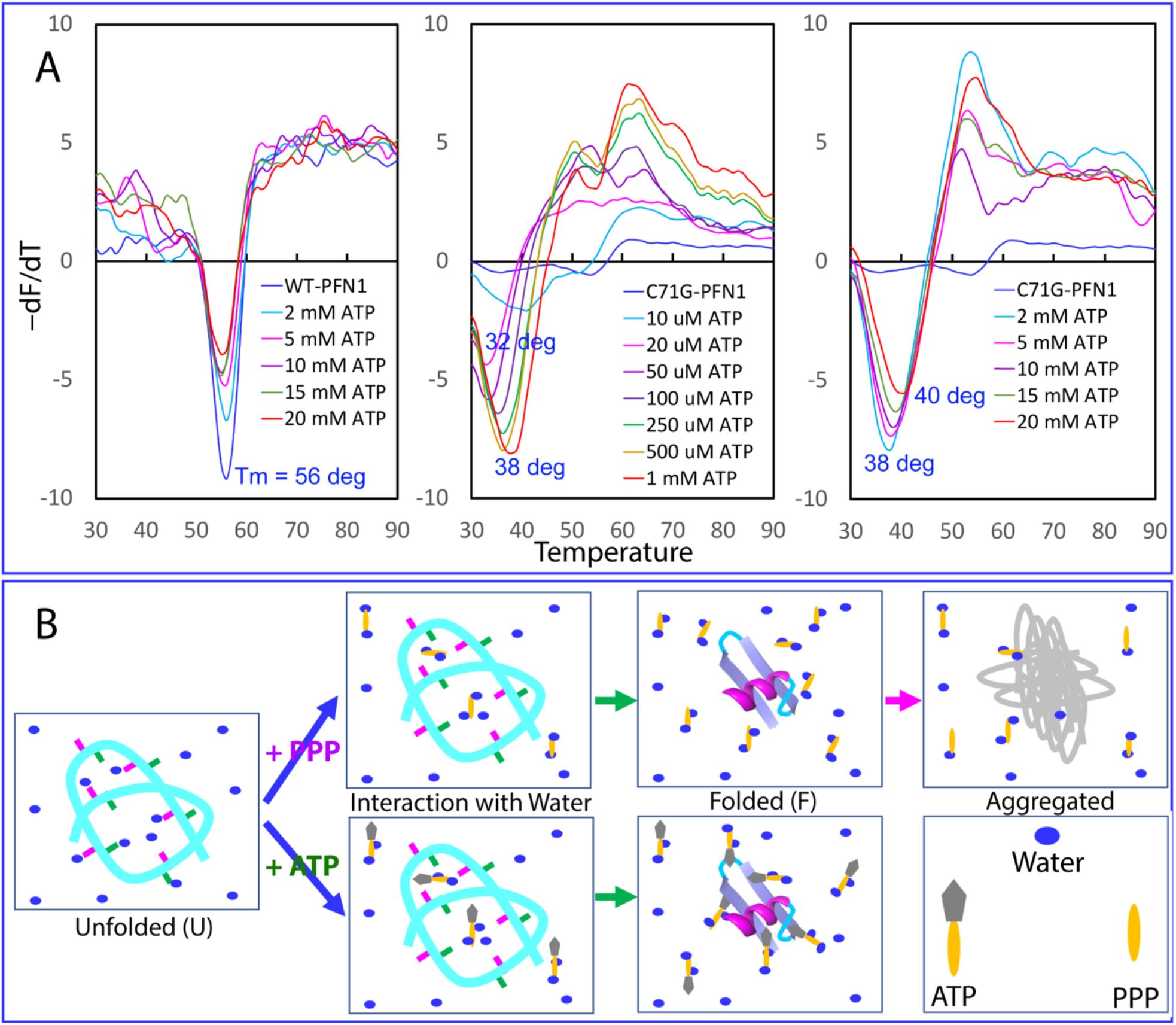
The speculative mechanism for ATP and PPP to induce protein folding. (A) ATP enhanced thermodynamic stability of C71G-hPFN1 but not WT-hPFN1 as measured by differential scanning fluorimetric (DSF). Melting curves of thermal unfolding of WT-hPFN1 and C71G-hPFN1 in the presence of ATP at different concentrations by plotting the first derivative of the fluorescence emission as a function of temperature (−dF/dT). Here Tm is represented as the lowest point of the curve. (B) A speculative model for protein folding induced by triphosphate (PPP) and ATP. For the unfolded state (U) of a protein, many backbone atoms are hydrogen-bonded with water molecules. Both triphosphate (PPP) and ATP can use their triphosphate chain to effectively attract water molecules out from hydrogen-bonding with the backbone atoms, thus favoring the formation of the folded state (F). However, due to the electrostatic screening effects produced by highly charged triphosphate, a protein exemplified by C71G-hPFN1 with significant exposure of hydrophobic patches due to the defects in tertiary packing will become severely aggregated. By contrast, ATP can use its aromatic base ring to dynamically interact with the exposed hydrophobic patches, which consequently not only functions to prevent aggregation, but also to increase the thermodynamic stability.

Previously our NMR studies suggest that ATP indeed has the high capacity in mediating the hydration of proteins even without needing of strong and specific binding (17,19,20,23,50). Recent NMR and MD simulation results also indicated that in general for proteins without the ability to bind nucleic acids, ATP only has very weak and non-specific effect on folded proteins even at very high ATP/protein ratios, which, however, appears to be sufficient to alter protein hydration (51). In this context, ATP appears to induce folding of C71G-hPFN1 only by enhancing the intrinsic folding capacity encoded by the amino acid sequence but introducing no additional information to specify folding. The results in the accompanying manuscript (47) also unveiled that unlike zinc ion which induces folding of the completely unfolded nascent hSOD1 by introducing the information additional to its amino acid sequence, ATP induces the folding of the nascent hSOD1 only by enhancing the intrinsic folding capacity encoded in its amino acid sequence. Indeed, even at 1:400, ATP induced no folding of intrinsically-disordered prion-like and RGG-rich domains of FUS (19) as well as prion-like domain of TDP-43 (20).

Our current study thus implies that nature selects triphosphate as the key intermediate for prebiotic evolution also because of its extremely-high efficiency in inducing protein folding. However, because of being highly charged, triphosphate at high concentrations will unavoidably provoke significant screening effect, thus triggering aggregation for the unfolded or partially-unfolded states of proteins as exemplified by C71G-hPFN1 here and nascent hSOD1 (47) with the significant exposure of hydrophobic patches. This may also explain why in most modern cells, the concentrations of triphosphate become extremely low. Strikingly, however, it has been reported that in some single-cell organisms, inorganic polyphosphates can function as a primordial chaperone, but the underlying mechanism remains elusive (31,32). Here we propose that polyphosphates might share the same mechanism of triphosphate to generally enhance protein folding by enhancing the intrinsic folding capacity encoded by the protein sequences. In this context, we further propose that ATP might also play a central role in solving protein folding and aggregation problems in primitive cells which lacked any modern ATP-energy-dependent machineries. Therefore, ATP appears to play the irreplaceable roles in the Origin of Life. Nevertheless, even in modern cells, ATP appears to still operate at fundamental levels to energy-dependently control protein homeostasis together with the energy-dependent chaperone and disaggregase machineries. This may rationalize the long-standing observation that the risk of neurodegenerative diseases dramatically increases with being aged, which is characterized by the constant reduction of ATP concentrations. Finally, our results may also offer a novel principle to engineer molecules like ATP capable of efficiently inducing folding, inhibiting aggregation and increasing stability to treat aggregation-associated ageing and diseases, as well as for various other applications.

## Materials and Methods

### Chemicals

ATP, ADP, AMP, sodium tripolyphosphate (PPP), sodium pyrophosphate (PP), ATPP, AMP-PCP, AMP-PNP, Adenosine, sodium phosphate, sodium chloride and trimethylamine N-oxide (TMAO) were all purchased from Sigma-Aldrich as previously reported (15,18-23). The fluorescent dye SYPRO Orange (S5692–50UL) was purchased from Sigma-Aldrich (21,23,43).

In our previous study (27), we have systematically evaluated the effects of pH and salts including NH_4_Cl, NaCl, KCl, CaCl_2_ and MgCl_2_ on the conformations and aggregation of C71G-hPFN1. The results indicated that: 1) at pH above 6.5, considerable aggregation occurred; 2) no ion had any detectable ability to induce folding but all triggered precipitation at high concentrations. Therefore in the present study, protein samples, as well as the list of compounds (Fig. 1) were all prepared in 1 mM sodium phosphate buffer containing 2 mM DTT with the final pH adjusted to 6.0 with very diluted NaOH or HCl.

### Preparation of WT- and C71G-hPFN1 proteins

The expression and purification of WT-hPFN1 and C71G-hPFN1, as well as removal of His-tag followed the protocol we previously described (27). To generate isotope-labeled proteins for NMR studies, the bacteria were grown in M9 medium with addition of (^15^NH_4_)_2_SO_4_ and ^13^-Glucose for ^15^N-/^13^C-labeling (27). The protein concentrations were determined by the UV spectroscopic method in the presence of 8 M urea, under which the extinct coefficient at 280 nm of a protein can be calculated by adding up the contribution of Trp, Tyr, and Cys residues (18-23,52,53).

### NMR titrations

All NMR experiments were acquired at 25 °C on an 800 MHz Bruker Avance spectrometer equipped with pulse field gradient units and a shielded cryoprobe as described previously (18-23,27,54,55). To conduct NMR titrations of ATP, ADP, AMP, ATPP, AMP-PCP, AMP-PNP, Adenosine, PPP, PP, sodium phosphate, sodium chloride and TMAO, two-dimensional ^1^H-^15^N NMR HSQC spectra were collected on the ^15^N-labeled C71G-hPFN1 at a protein concentration of 50 μM in 1 mM sodium phosphate buffer containing 2 mM DTT (with the final pH adjusted to 6.0) at 25 °C in the presence of ATP, ADP and AMP at molar ratios of 1:0.25, 1:0.5, 1:1, 1:2, 1:4, 1:6, 1:8, 1:10, 1:15, 1:20, 1:40, 1:100, 1:200, 1:300 and 1:400. For Adenosine, the ratios are 1:0.25, 1:0.5, 1:1, 1:2, 1:4, 1:6, 1:8, 1:10, 1:15, 1:20, 1:40, 1:100. For PPP and ATPP, the ratios are 1:0.25, 1:0.5, 1:1, 1:2, 1:4 where the sample started to precipitate. For PP, AMP-PCP and AMP-PNP, the ratios are 1:0.25, 1:0.5, 1:1, 1:2, 1:4, 1:6, 1:8, 1:10 where the sample started to precipitate. For sodium phosphate, the ratios are 1:1, 1:2, 1:10, 1:20, 1:40, 1:100 where the sample started to precipitate. For sodium chloride, the ratios are 1:1, 1:10, 1:20, 1:40, 1:100 and 1:200 where the sample started to precipitate.

NMR spectra were processed with NMR Pipe (56) and analyzed with NMR View (57). The molar ionic strength of the sodium salts were calculated as previously described (15,19,39).

### NMR sequential assignments

To achieve sequential assignments of both WT-hPFN1 and C71G-hPFN1, triple resonance NMR spectra HN(CO)CACB and CBCA(CO)NH were collected on the ^15^N-/^13^C-double labeled WT-hPFN1 or C71G-hPFN1 sample while HSQC-TOCSY and HSQC-NOESY were collected on the ^15^N-labeled WT-hPFN1 or C71G-hPFN1 sample. NMR ^1^H chemical shifts were referenced to external DSS at 0.0 ppm (18-23,27,55).

### PGF Diffusion Measurement

PGF-NMR experiments were run on an 800 MHz Bruker Avance spectrometer at 25 ºC. The different protein samples were prepared in D_2_O at protein concentration of 50 μM in 1 mM sodium phosphate buffer (pD 6.0). The experiments were performed using the Bruker pulse sequence and the Bruker macro diffusion ordered spectroscopy (DOSY) (30,38). Typically 16 values of gradient strength were used in the range 0 to 32 G/cm, with PFG duration of 2 ms, and diffusion time of 150 ms. The self-diffusion coefficients (*D*_*s*_) were calculated using the Bruker DOSY analysis program. Each sample was run in triplicate and *Ds* values were averaged over the three experiments. The resulting decay curves were fitted and *Ds* values were calculated with the equation below:

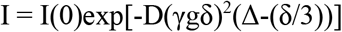

Where I(0) is 1.002, γ is 4.258×10^3^ Hz/G, δ is 4.000 ms, and Δ is 150 ms.

### Quantification of Exchange Kinetics

Longitudinal magnetization transfer due to chemical exchange is the basis for the appearance of exchange cross peaks in nuclear Overhauser effect spectroscopy (NOESY) spectra. The evolution of longitudinal magnetization is described by (29):

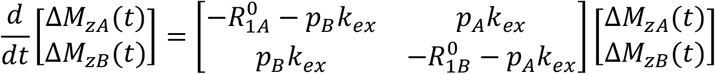

In which 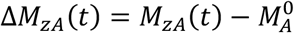 and 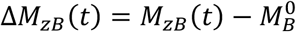. The solution of this equation is:

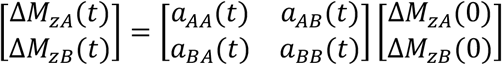

In which

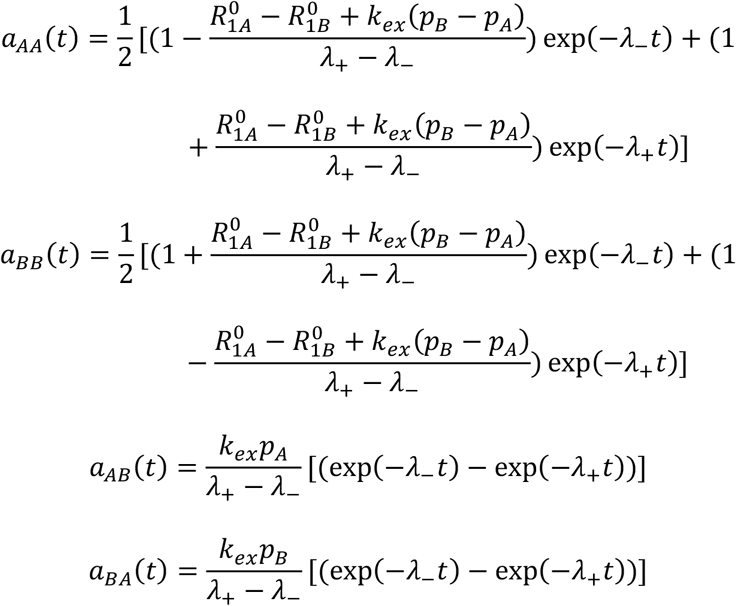

and

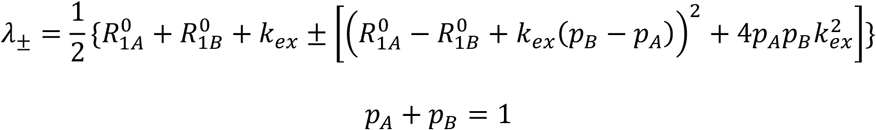

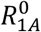 and 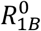 were measured by separate experiments. As a consequence, here *k*_*ex*_, *p*_*A*_, and *p*_*B*_ could

be derived as previously described (29,30) and presented in Table S1.

### NMR ^15^N backbone dynamics on ps-ns time scale

^15^N backbone T1 and T1ρ relaxation times and {^1^H}-^15^N steady state NOE intensities were collected on the ^15^N-labeled WT- and C71G-hPFN1 at 25 °C in 1 mM phosphate buffer (pH 6.0) on an Avance 800 MHz Bruker spectrometer with both an actively shielded cryoprobe and pulse field gradient units (34-37). Relaxation time T1 was determined by collecting 7 points with delays of 10, 160, 400, 500, 640, 800 and 1000 ms using a recycle delay of 1 s, with a repeat at 400 ms. Relaxation time T1ρ was measured by collecting 8 points with delays of 1, 40, 80, 120, 160, 200, 240 and 280 ms, with a repeat at 120 ms. {^1^H}-^15^N steady-state NOEs were obtained by recording spectra with and without ^1^H presaturation, a duration of 3 s and a relaxation delay of 6 s at 800 MHz.

### Model-free analysis

NMR relaxation data were analyzed by “Model-Free” formalism with protein dynamics software DYNAMICS (34-37). Briefly, relaxation of protonated heteronuclei is dominated by the dipolar interaction with the directly attached ^1^H spin and by the chemical shift anisotropy mechanism. Relaxation parameters are given by:

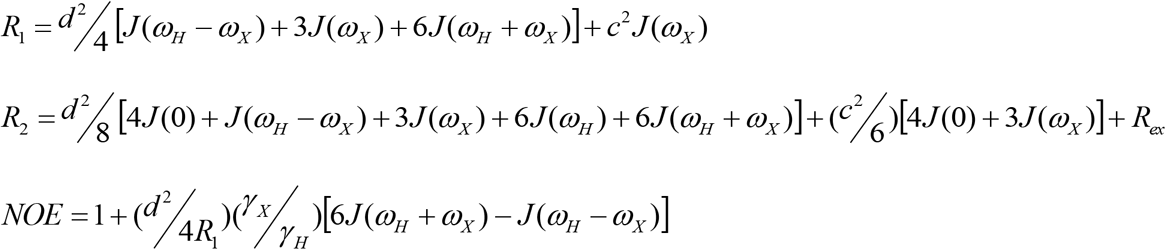

In which,

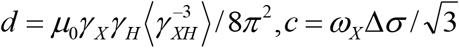, *μ*_0_ is the permeability of free space; *h* is Planck’s constant; *γ*_*X*_, *γ*_*H*_ the gyromagnetic ratios of ^1^H and the X spin (X=^13^C or ^15^N) respectively; *γ* is the X-H bond length; *ω*_*H*_ and *ω*_*X*_ are the Larmor frequencies of ^1^H and X spins, respectively; and Δ*σ* is the chemical shift anisotropy of the X spin.

The Model-Free formalism determines the amplitudes and time scales of the intramolecular motions by modeling the spectral density function, *J*(*ω*), as

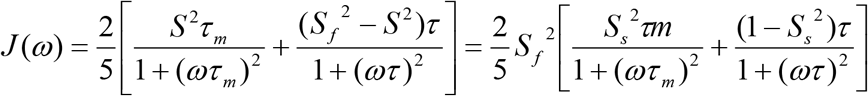

In which, 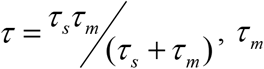 is the isotropic rotational correlation time of the molecule, *τ*_*s*_is the effective correlation time for internal motions, *S* ^2^ = *S*_*f*_ ^2^*S*_*s*_ ^2^ is the square of the generalized order parameter characterizing the amplitude of the internal motions, and *S*_*f*_ ^2^ and *S*_*s*_ ^2^ are the squares of the order parameters for the internal motions on the fast and slow time scales, respectively.

In order to allow for diverse protein dynamics, several forms of the spectral density function, based on various models of the local motion, were utilized, which include the original Lipari-Szabo approach, assuming fast local motion characterized by the parameters *S*^2^ and *τ*_*loc*_; extended model-free treatment, including both fast (*S*_*fast*_ ^2^, *τ*_*fast*_) and slow (*S*_*slow*_ ^2^, *τ*_*slow*_) reorientations for the NH bond (*τ*_*fast*_ ≪ *τ*_*slow*_ < *τ*_*c*_); and could also allow for slow, milli- to microsecond dynamics resulting in a conformational exchange contribution, *R*_*ex*_, to the linewidth. In DYNAMICS, there are eight models for local motions and each residue is fitted with different models. Subsequently goodness of fit will be checked and the best-fitted model will be selected.

The relaxation data of WT-hPFN1 and the folded state of C71G-hPFN1 were analyzed with the previously published X-ray structure (pdb ID of 2PAV) (24) by isotropic, axially-symmetric and fully anisotropic models for the overall motion and the results were tested and then compared. According to the illustration of ROTDIF, isotropic model was finally selected for both WT-hPFN1 and the folded state of C71G-hPFN1 because of smallest Ch^2^/df value. For WT-hPFN1, τc = 7.5 ns and Dx = Dy = Dz = 1.852 E ± 07 s^-1^; for the folded state of C71G-hPFN1, τc = 7.8 ns and Dx = Dy = Dz = 2.094 E ± 07 s_-1_.

### Determination of thermodynamic stability by DSF

As ATP triggers significant noise in circular dichroism (CD) measurement, as well as the quenching effect of the intrinsic Trp fluorescence as we previously reported (19,21-23), therefore we were unable to use CD and fluorescence spectroscopy to determine the thermodynamic stability of C71G-hPFN1 in the presence of ATP and PPP by performing chemical and thermal unfolding as usually conducted (37). Here we thus measured the thermodynamic stability of the WT- and C71G-hPFN1 by differential scanning fluorimetric (DSF) method (43), as we previously used to characterize the effect of ATP on other proteins (21-23). DSF was used to determine the thermodynamic stability of WT-hPFN1 and C71G-hPFN1 at protein concentrations of 10 μM in 1 mM sodium phosphate buffer containing 2 mM DTT (pH 6.0) in the presence of ATP at different concentrations. DSF experiments were performed using the CFX384 Touch™ Real-Time PCR Detection System from BIO-RAD, following the SYBR green melting protocol to obtain Tm value. Briefly, in a single well of a 384-well PCR plate, a 10-μL reaction solution was placed, which contains the proteins at 10 μM, ATP at different concentrations, and 10× SYPRO Orange. Plates were sealed with a quantitative PCR adhesive optical seal sheet (Microseal ‘B’ Adhesive Sealing Films, BIO-RAD) and then spun at 1000 rpm for 1 min to remove bubbles. The program in Real-Time PCR instrument was set to SYBR green and ran the temperature scan from 25 °C to 95 °C with the increment of 1 °C/min. Upon completion, the obtained thermal unfolding curves were displayed as the first derivatives (dF/dT) by the reverse transcriptase PCR software Bio-Rad CFX Manager 3.0.

## Acknowledgments

This study is supported by Ministry of Education of Singapore MOE Tier 1 A-8000711-00-00 Grant to Jianxing Song.

## Author contributions

J.S. conceived and designed the research; J.K., L.-Z.L., and J.S. performed research; L.Z.L., J.K., and J.S. analyzed data; and J.S. wrote the paper.

